# Combinatorial DNA-PAINT

**DOI:** 10.64898/2026.01.23.701249

**Authors:** Philipp R. Steen, Luciano A. Masullo, Isabelle Pachmayr, Rafal Kowalewski, Larissa Heinze, Clemens Steinek, Jisoo Kwon, Monique Honsa, Susanne C. M. Reinhardt, Heinrich Grabmayr, Ralf Jungmann

## Abstract

State-of-the-art super-resolution microscopy enables nanometer-resolution imaging of proteins, but its multiplexing capacity has been fundamentally constrained. Here, we present Combi-PAINT, a combinatorial DNA barcoding strategy based on DNA-PAINT, that facilitates superlinear scaling of multiplexing with the number of imaging rounds. Instead of assigning one unique docking strand per target, Combi-PAINT encodes targets as combinations of orthogonal sequences, allowing nonlinear scaling of target number with imaging rounds. Using just six imaging rounds, we resolve 41 targets in a field-of-view of 100 x 100 μm^2^ with ∼2.5 nm localization precision and ∼90% decoding accuracy in under 30 minutes, representing the fastest sub-10 nm super-resolution microscopy acquisition to date. We benchmark decoding fidelity and demonstrate robust *in situ* performance in mammalian cells, achieving 97% decoding accuracy. Combi-PAINT is compatible with existing DNA-PAINT workflows and speed-enhancing techniques, offering a scalable, accessible platform for high-content, single-molecule spatial proteomics.

## Introduction

A comprehensive, mechanistic understanding of biological systems relies on the ability to precisely map the spatial distribution, abundance, and molecular interactions of proteins. Proteins orchestrate virtually every biological process, from signal transduction and immune recognition to cell division and tissue organization. These processes are often governed by the nanoscale spatial arrangement and interplay of large numbers of distinct protein species. Importantly, many regulatory mechanisms – such as receptor clustering or the assembly of signaling hubs – occur at spatial scales well below the diffraction limit of light. Therefore, to decode how cells function, adapt, or malfunction in disease, it is essential to measure protein positions and stoichiometries at single-molecule resolution and across highly multiplexed molecular contexts.

Mass spectrometry can quantify thousands of protein species at single-cell resolution^1^, and recent advances achieve subcellular resolutions down to hundreds of nanometers^2-4^. However, only super-resolution microscopy^5^ can ultimately achieve the spatial precision and molecular specificity necessary for resolving individual biomolecules *in situ*^6-9^. Multiplexing capabilities in traditional super-resolution microscopy are often limited by spectral overlap and the finite number of distinguishable fluorophores. DNA-PAINT^10,11^ overcomes these constraints by leveraging the transient hybridization of short, dye-labeled “imager” strands to complementary, target-bound “docking” strands, producing single-molecule blinking. This method enables high degrees of multiplexing through Exchange-PAINT^12^, in which orthogonal DNA sequences are imaged sequentially. Consequently, DNA-PAINT can visualize dozens of distinct protein species within the same sample.

Exchange-PAINT and related sequential imaging approaches^13,14^ theoretically allow for proteome-scale multiplexing by imaging orthogonal docking strands in separate rounds. However, a fundamental limitation remains: the total acquisition time scales linearly with the number of targets, creating a major bottleneck for high-throughput and high-content imaging applications. For instance, imaging 50 targets requires 50 separate imaging rounds, each involving buffer exchanges, washing steps, and drift corrections. This severely limits throughput, especially for large fields of view or large-scale spatial proteomics studies across many samples or conditions. To address this challenge, combinatorial barcoding strategies have emerged as powerful approaches to decouple multiplexing capacity from the number of imaging rounds. These approaches assign unique combinations of labels to each target, enabling superlinear scaling of distinguishable identities relative to the number of labels or rounds. While such strategies have been successfully implemented in RNA imaging (MERFISH^15^ and seqFISH^16^) and diffraction-limited protein imaging at the tissue level^17^, combinatorial multiplexing for single-molecule protein imaging with nanometer precision has not yet been achieved.

Here, we introduce Combi-PAINT, a method that implements combinatorial barcoding directly within the DNA-PAINT framework. Instead of labeling each target with a single unique docking sequence, Combi-PAINT encodes targets with specific combinations of docking “base sequence” motifs on the same docking strand. Each imaging round reads out the presence of a specific base sequence motif, and target identities are reconstructed from the presence of specific signals across rounds. This strategy enables a nonlinear increase in multiplexing capacity, significantly reducing the number of imaging rounds required.

## Results

### Combinatorial DNA-PAINT

In conventional sequential imaging approaches such as Exchange-PAINT^12^, each molecular target is labeled with a unique DNA-PAINT docking sequence – hereafter referred to as a base sequence (**Fig. 1a**). These targets are imaged sequentially, with one imaging round per base sequence (**Fig. 1b**). Buffer exchanges between rounds ensure unambiguous readout (**Fig. 1c**). While this approach enables high-resolution imaging of multiple targets, its scalability is fundamentally constrained since the total acquisition time increases linearly with the number of target species.

**Figure 1:**
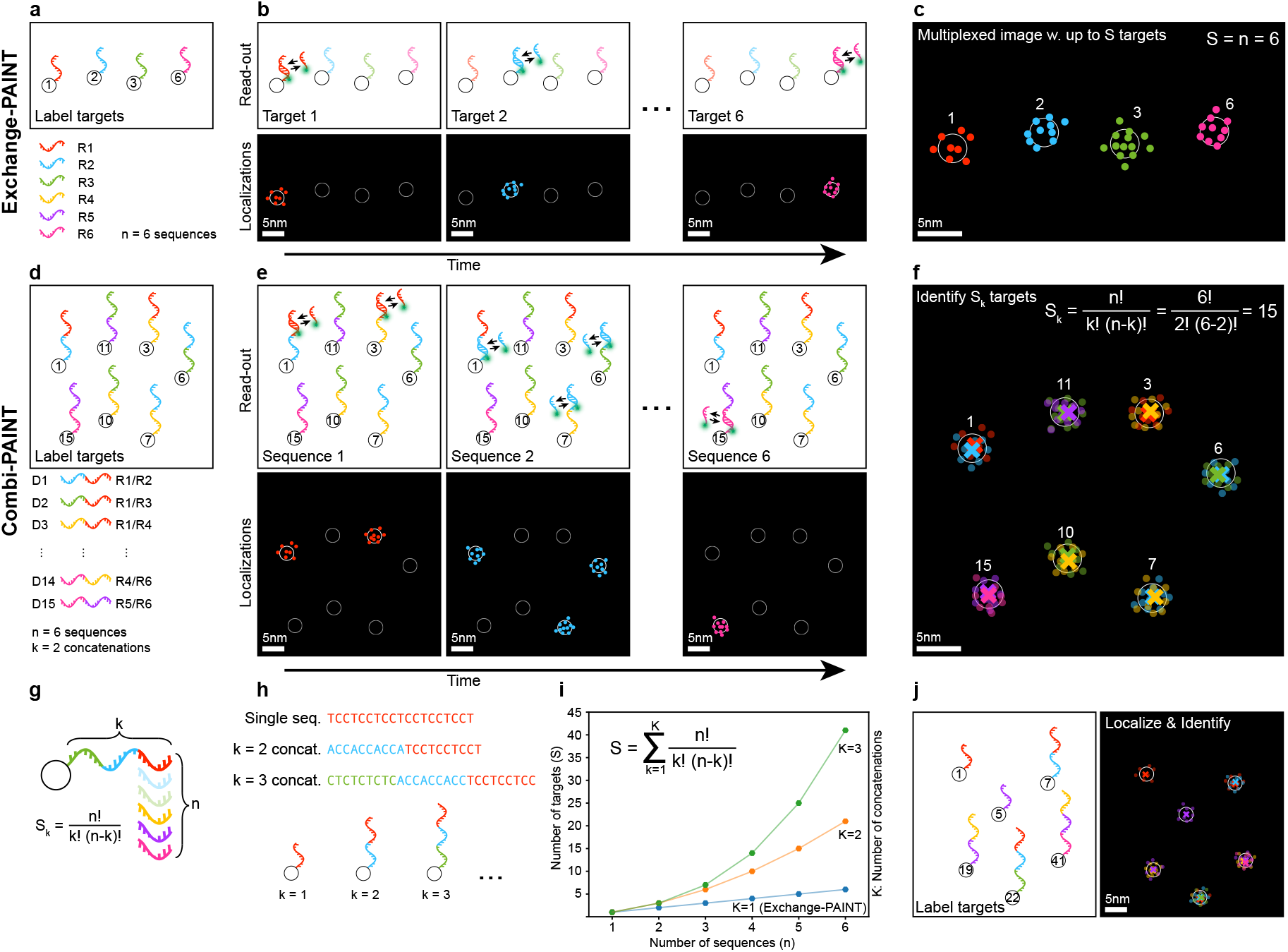
Enhanced multiplexing via combinatorial DNA-PAINT. **a**, conventional labeling assigns one docking sequence to each target. **b**, the targets are read-out sequentially via Exchange-PAINT or related methods, gathering localizations from one target species per imaging round. **c**, the degree of multiplexing (S) achieved is equal to the number of sequences used (n), which is equal to the number of imaging rounds performed. **d**, in Combi-PAINT, each target is labeled with a sequence containing one or several (k), in this case two, out of n base sequence docking motifs. **e**, sequential imaging gathers localizations belonging to multiple target species labeled with the same base sequence segments. **f**, overlaying all localizations and assigning target identities based on localization content, achieves combinatorial imaging with S = 15 encodable target species using n = 6 base sequences and imaging rounds. **g**, the degree of multiplexing S is determined by the number of concatenations k and the number of available base sequences n. **h**, Single (k = 1), double (k = 2) and triple (k = 3) docking motif sequences may be used in conjunction, leading to a new scaling shown in (**i**): using k = 1, 2, 3 and n = 6 sequences yields potentially 41 target species to be imaged in 6 imaging rounds, as illustrated in (**j**).

Combi-PAINT overcomes this limitation by introducing a combinatorial encoding strategy. Instead of assigning a single base sequence to each target, Combi-PAINT labels targets with specific combinations of multiple base sequences (**Fig. 1d**). Each imaging round then detects the presence of one base sequence across the entire sample (**Fig. 1e**), and the identity of each labeled target is reconstructed by analyzing the pattern of detected signals across all imaging rounds (**Fig. 1f**). This strategy enables nonlinear scaling of multiplexing capacity with the number of imaging rounds, dramatically improving throughput.

Formally, if *n* base sequences (and corresponding imaging rounds) are available, and each target is labeled with a specific combination of *k* sequences (**Fig. 1g, h**), the number of uniquely encodable targets *S*_*k*_ is given by the binomial coefficient:

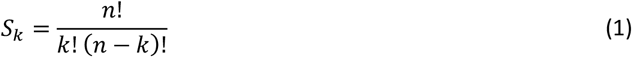

The total number of distinct encodable target species S that may be labeled with combinations of k = 1 to K different base sequences (see **Fig. 1i**) is the sum of S_k_ over all k:

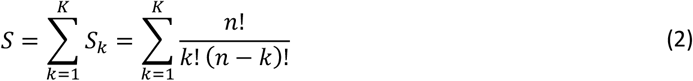

For example, using *n = 6* base sequences and allowing combinations of size *k = 1*, 2, *3* (which we refer to as R, D, and T sequences), a total of 41 unique barcode identities can be generated and read out in just 6 imaging rounds (**Fig. 1j**). Crucially, this combinatorial encoding allows simultaneous detection of many targets without the need to linearly increase imaging time, offering a scalable solution for high-multiplex, high-throughput applications. We quantified this scaling by comparing the relative acquisition time per target (**Supplementary Note 1, Supplementary Figure 1**). For 15 targets (according to **Eq. 1**), the time required per target is less than half that of conventional individual DNA-PAINT measurements.

### Benchmarking using DNA origami

To experimentally validate Combi-PAINT, we first benchmarked its performance in a controlled *in vitro* setting using well-defined DNA origami structures. As base sequences, we employed speed-optimized R strands^18^ to construct combinatorial barcodes comprising either two (D sequences, *k = 2*) or three (T sequences, *k = 3*) concatenated R strands. For example, the D1 sequence consists of R1 and R2, D2 of R1 and R3, and T1 of R1, R2, and R3 (see **Supplementary Table 1** for full sequence designs).

We designed rectangular DNA origami structures^19^ containing 12 identical Combi-PAINT docking sites spaced 20 nm apart^11^ (**Fig. 2a**). After validating R-, D-, and T-sequence DNA origami in separate experiments, we pooled all 41 barcode species – containing 6 R (single), 15 D (double), and 20 T (triple) sequences – and imaged them in six Exchange-PAINT rounds. The average localization precision achieved was 2.5 ± 0.3 nm. Representative origami structures from each barcode class are shown in **Fig. 2b**, and channel-by-channel localization maps for a subset of structures are displayed in **Fig. 2c**.

**Figure 2:**
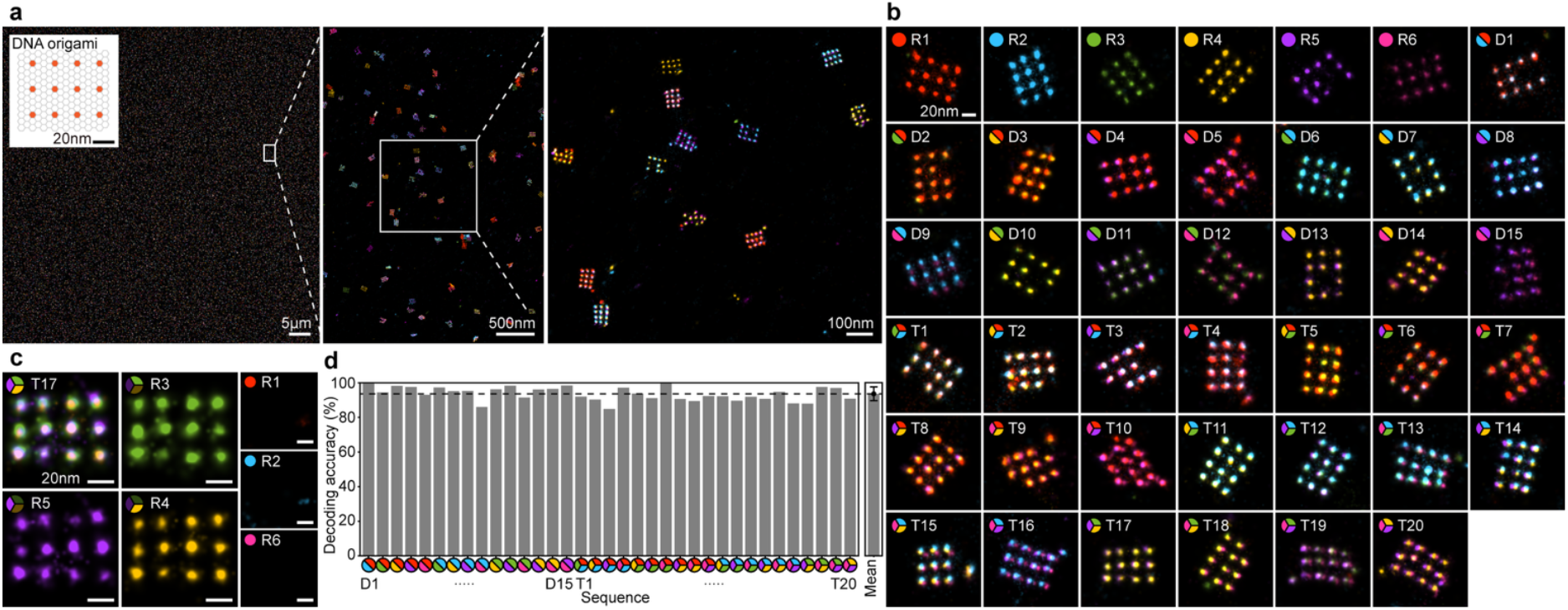
41-plex Combi-PAINT in DNA origami. **a**, DNA-PAINT FOV of 75 x 75 μm^2^ containing 41 different DNA origami species (insert shows DNA origami design) imaged in 3 h and 20 min at a localization precision σ_NeNA_ of approx. 2.5 nm. **b**, 41 DNA origami species presenting k=1 (R1 - R6), k=2 (D1 - D15) or k=3 (T1 - T20) docking motifs are present and resolved. **c**, displaying the three underlying imaging rounds shows localizations present at all docking sites, while very few localizations are present in the other three channels. **d**, the mean decoding accuracy (portion of correctly identified docking site identities) is 94 ± 4%.

The identity of each DNA origami (R1-R6, D1-D15, or T1-T20) was retrieved by analyzing all its localizations pooled together (see **Methods** for details). Next, the Combi-PAINT analysis workflow (**Extended Data Figure 1**) was applied: (1) Localizations from all channels (**Extended Data Figure 1a**) were combined into a single dataset while retaining their channel identities. (2) A gradient ascent-based radial grouping algorithm^7^ was applied to identify single docking sites (**Extended Data Figure 1b**). (3) Docking sites were filtered according to their binding kinetics to remove unspecific “sticking” events (see **Methods, Supplementary Note 2**, and **Supplementary Figure 2** for more details). (4) Finally, the channel identities of all localizations per binding site were used to determine the combinatorial identity (**Extended Data Figure 1c, d**).

We then quantified the decoding accuracy (DA), defined as the fraction of binding sites assigned the correct barcode, compared to the ground truth determined previously. For D sequences, we obtained a DA of 96 ± 3%, while T sequences yielded 92 ± 4%, resulting in an overall mean DA of 94 ± 4% across all 41 barcodes (**Fig. 2d**). To assess the upper limits of acquisition speed, we repeated the 41-plex imaging using 4-minute imaging rounds, resulting in a total imaging time of 24 minutes. Despite the shorter acquisition, decoding yielded a DA of 89 % (CI: 74 - 100 %), corresponding to an effective imaging rate of 30 seconds per target across a field of view (FOV) of 100 × 100 µm^2^. This acquisition speed represents an order-of-magnitude improvement over previously reported DNA-PAINT methods at comparable resolution and multiplexing levels.

### Evaluation on surface receptors

While Combi-PAINT exhibits high decoding accuracy in well-defined DNA origami structures, applying it in cellular environments introduces new challenges, such as increased spatial crowding, background signal, and potentially decreased docking site accessibility. We thus evaluated the performance of Combi-PAINT *in situ*, targeting cell surface receptors.

We first generated anti-GFP nanobodies labeled with each D sequence and tested them targeting CD86-GFP in transiently transfected CHO-K1 cells (**Fig. 3a**). Each D sequence (D1-D15) was imaged in a separate experiment with the two imaging channels corresponding to its base R components (e.g., D3 = R1 + R4). The resulting localizations showed well-defined clusters (**Fig. 3b**), with an average localization precision across all measurements of 2.9 ± 0.3 nm.

**Figure 3:**
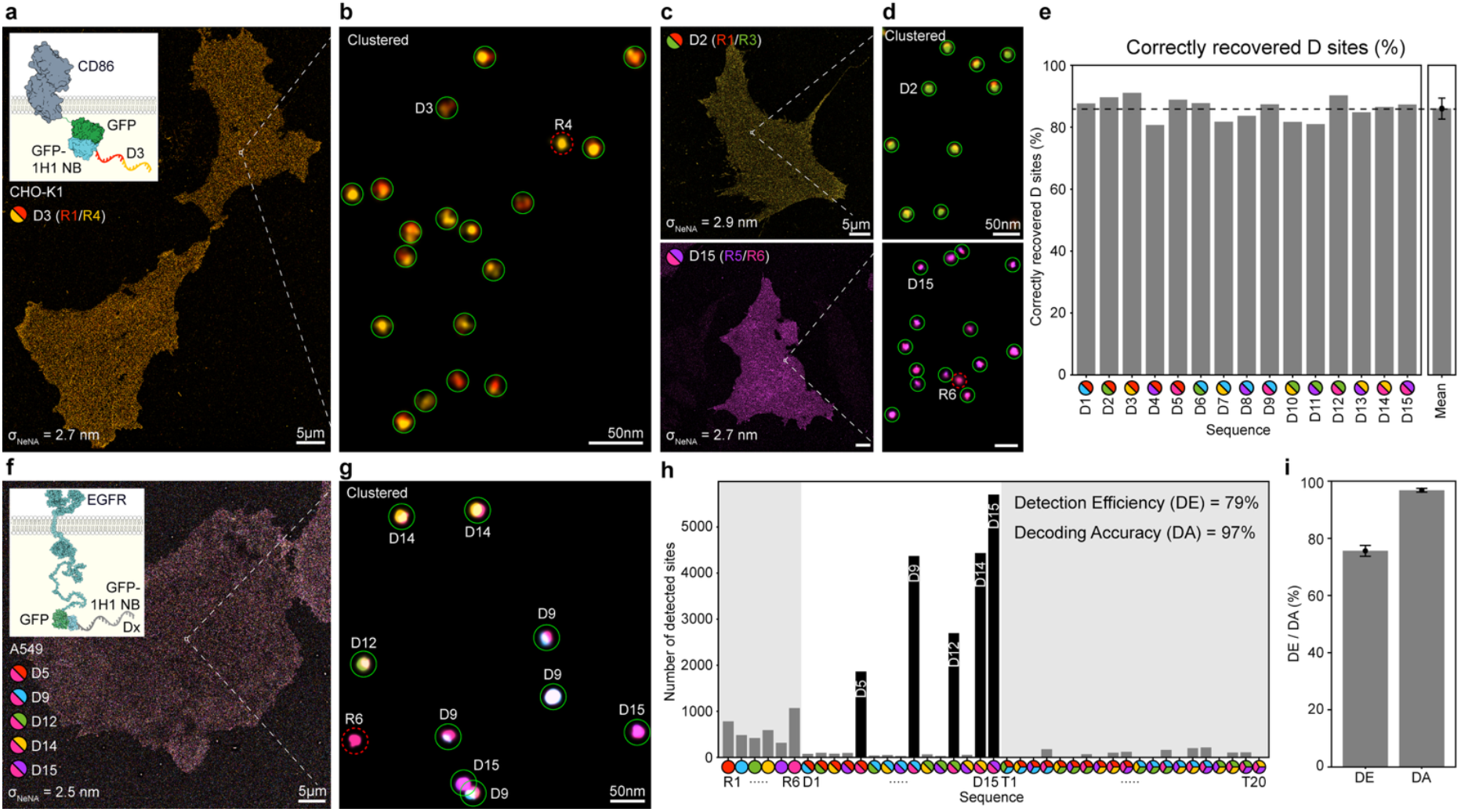
Combi-PAINT in cells. **a**, CHO-K1 cells transiently transfected with CD86-GFP and labeled with D3 anti-GFP nb imaged in R1* and R4* channels. **b**, Combi-PAINT analysis yields mostly correctly identified D3 sites (indicated by green circles) and some incorrectly identified sites (R4, red dotted circle). **c**, such experiments were repeated for all D sequences; D2 and D15 are shown. **d**, results are consistent across D sequences with occasional instances where D sequences are incorrectly identified as R sequences. **e**, the percentage of correctly recovered D sites is 86 ± 3 %. **f**, to quantify D sequence decoding accuracies, A549 cells expressing EGFR-GFP were labeled with five D sequences (D5, D9, D12, D14, D15) in roughly equal proportions and imaged in all R1*-R6* channels. **g**, sites are mostly correctly identified. **h**, the detection efficiency was 79% while the decoding accuracy was 97%. **i**, for N=4 measurements using different combinations of D sequences, the overall DE is 76 ± 2% and overall DA is 97 ± 1%.

Applying Combi-PAINT analysis, cells labeled with D3, for example, yielded mostly correctly recovered targets (green circles in **Fig. 3b**), with a small number of D→R-type misassignments (red dotted circle in **Fig. 3b**). Additional experiments using other D sequences (e.g., D2, D15) showed similar results (**Fig. 3c, d**). Across all D-sequence experiments, the mean percentage of correctly recovered D sites was 86 ± 3% (**Fig. 3e**).

To evaluate decoding performance in a multiplexed context, we labeled A549 cells expressing EGFR-GFP with a pool of five D sequences (D5, D9, D12, D14, and D15), each targeting EGFR via anti-GFP nanobodies. Cells were imaged across all six R channels. A representative FOV showed correct identification of all five target species and minimal false-positive detections (**Fig. 3f, g**).

For this experiment, the preliminary decoding accuracy over all 41 channels was 76%. However, if we restrict the coding space to D sequences only (and thus discard decoded T and R sequences), the decoding accuracy is increased to 97% (**Fig. 3h**) at the expense of a reduction in the detection efficiency to 79% (see **Methods** for details). We repeated this analysis using stably transfected CHO-K1 cells expressing CD86-GFP labeled with another subset of D sequences (D2, D8, D10, D14, D15), achieving a decoding accuracy of 97 ± 1% (and a detection efficiency of 76 ± 2%) across four independent experiments (**Fig. 3i**).

## Discussion

With Combi-PAINT, we introduce a new framework for high-throughput, multiplexed DNA-PAINT imaging that overcomes the traditional trade-off between resolution, multiplexing capability, and acquisition time. The core conceptual advance is the transformation of the functional relationship between the number of imaging rounds and the number of imaged targets through combinatorial barcoding. By assigning each target a unique combination of DNA sequences instead of a single one, the number of distinguishable targets increases nonlinearly with the number of imaging rounds.

Compared to existing sequential methods, Combi-PAINT eliminates the need for one-to-one target-to-round mapping. As a result, we demonstrate the fastest sub-10 nm super-resolution microscopy acquisition to date, resolving 41 targets in under 30 minutes over a 100 x 100 μm^2^ field of view with 2.5 nm localization precision. Importantly, this speed gain does not compromise accuracy: even in complex cellular environments, Combi-PAINT achieves 97% accuracy in decoding combinatorially labeled surface receptors.

As shown by our biological proof-of-principle experiments, Combi-PAINT operates robustly in complex cellular environments and can be readily applied to interrogate cellular systems relevant to e.g. cancer diagnostics and immunotherapy. Immediate applications include the nanoscale analysis of immune checkpoints and other immunological targets, where the precise spatial organization and molecular interplay of receptors and ligands critically influence signaling outcomes.

To fully unlock the potential of Combi-PAINT for broad biological discovery, an expanded repertoire of high-quality binders will be essential. Improving binder affinity and specificity will be crucial to reliably label and decode a wider range of molecular targets. Advances in *de novo* protein design^20^ could play a transformative role in this regard, enabling systematic access to previously intractable proteins and thereby further scaling the biological reach of Combi-PAINT.

Technically, Combi-PAINT is simple to implement: it requires no additional chemistry, enzymatic barcoding, or optical multiplexing, and builds directly on standard Exchange-PAINT workflows. It is also compatible with existing speed-enhancing methods such as simultaneous multi-color imaging^21^, which could reduce imaging rounds by a factor of two or even three. From a hardware point of view, Combi-PAINT is compatible with automation and integration into high-content screening pipelines^22^.

Finally, Combi-PAINT benefits directly from recent advances in DNA sequence design, specifically the use of left-handed DNA strands^23^ and the expansion of the speed-optimized sequences^24^ from 6 to 10. Therefore, Combi-PAINT could be conducted with 20 base sequences (10 speed-optimized right- and their left-handed counterparts). This would lead to up to 190-plex in 20 speed-optimized imaging rounds (∼12 hours of imaging time) at high decoding accuracy. Incorporating triple combinations (**Eq. 2**) would increase this number to >1300 available channels, making Combi-PAINT a method capable of imaging the whole surfaceome of the cell at single-protein resolution using a fluorescence microscope.

## Extended Data Figures

**Extended Data Figure 1.**
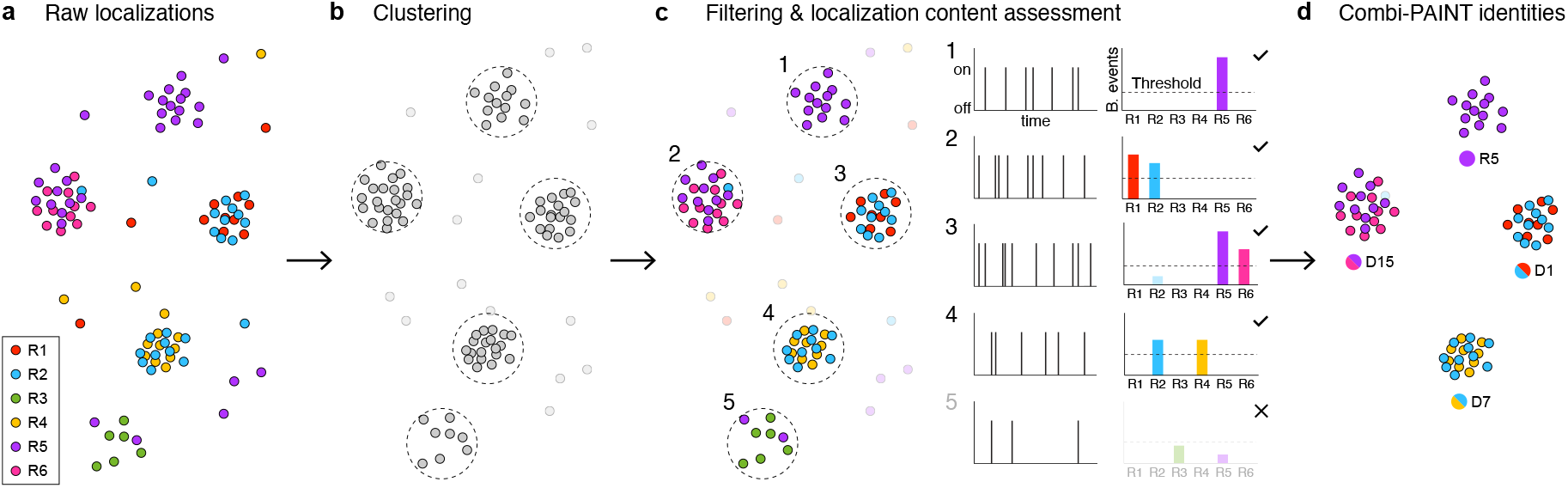
Combi-PAINT analysis workflow. **a**, Raw localizations from six imaging rounds (R1 to R6). **b**, Localizations are merged, and a clustering algorithm is applied to the merged dataset while imaging round information of each localization is preserved. **c**, Each cluster of localizations is analyzed, the time trace of binding events is extracted, and the combinatorial identity of the binding site is decoded from the localization content. Furthermore, clusters are also filtered according to the number of binding events and binding kinetics. **d**, The final Combi-PAINT image is rendered by merging the multi-base-channel localizations into the corresponding decoded combinatorial channel.

## METHODS

### Materials

A549 Cells GFP-EGFR (CLL1141-1VL, Sigma-Aldrich) and CHO-K1 cells (CCL-61) were obtained from ATCC. DNA oligonucleotides modified with C3-azide, Cy3B, and DBCO were ordered from Metabion and MWG Eurofins. DNA oligonucleotides modified with Gly-Gly-Gly were ordered from Biomers. Magnesium chloride (1 M; AM9530G), sodium chloride (5 M; AM9760G), potassium chloride (2 M; AM9640G), calcium chloride (1 M, 15445389), ultrapure water (10977-035), Tris (1 M, pH 8; AM9855G), EDTA (0.5 M, pH 8.0; AM9260G), 1× PBS (pH 7.2; 20012-019), 10× PBS (70011051), FBS (10500-064), 0.05 % trypsin–EDTA (25300-054), EZ-Link Maleimide-PEG4-DBCO (cat: C20041), and Salmon Sperm DNA (15632011) were purchased from Thermo Fisher Scientific. BSA (A4503-10G), TritonX-100 (93443), and Millipore Millex 33mm MCE 0.22 µm sterile filter (SLGS033) were ordered from Sigma-Aldrich. Amicon Ultra Centrifugal Filters with a 100-kDa molecular weight cutoff (MWCO, UFC510096) and those with a 10-kDa MWCO (UFC201024) were ordered from Merck Millipore. Ammonium chloride (K298.1) was purchased from Carl Roth. Sodium hydroxide (31627.290) was purchased from VWR. Methanol-free paraformaldehyde (15710) was obtained from Electron Microscopy Sciences. Glutaraldehyde (23115.01) was purchased from SERVA. Tween-20 (P9416-50ML), glycerol (65516-500ml), methanol (32213-2.5L), protocatechuate 3,4-dioxygenase pseudomonas (PCD; P8279), 3,4-dihydroxybenzoic acid (PCA; 37580-25G-F), (±)-6-hydroxy-2,5,7,8-tetra-methylchromane-2-carboxylic acid (trolox; 238813-5G), sodium azide (769320) were obtained from Sigma-Aldrich. 90-nm gold nanoparticles (G-90-100) were ordered from Cytodiagnostics. Anion exchange column RESOURCE Q (17117701) was obtained from Cytiva. Nanobodies against GFP (clone 1H1, N0305) with a single ectopic cysteine at the C-terminus for site-specific conjugation were purchased from Nanotag Biotechnologies. DBCO-PEG4-Maleimide (CLK-A108P) and DBCO-PEG4-Gly-Gly-Gly linker (Vectorlabs, cat. no. CCT-1552) were purchased from Jena Bioscience. Microscope slides were obtained from ibidi (μ-Slides with 8 wells with a glass bottom, 80801).

### Buffers

- Buffer A: 10 mM Tris-HCl pH 8, 100 mM NaCl and 0.05% Tween-20; pH 8
- Buffer B: 10 mM MgCl_2_, 5 mM Tris-HCL pH 8, 1 mM EDTA and 0.05% Tween-20; pH 8
- Buffer C: 1× PBS, 0.1 mM EDTA and 500 mM NaCl, 0.05%Tween; pH 7.4
- Folding buffer: 10 mM Tris, 1 mM EDTA, 12.5 mM MgCl_2_; pH 8
- FoB5 buffer: 5 mM Tris, 1 mM EDTA, 5 mM NaCl, 5 mM MgCl_2_; pH 8
- Blocking buffer: 1× PBS, 1 mM EDTA, 0.02% Tween-20, 0.05% NaN_3_, 2% BSA, 0.05 mg ml^−1^ sheared salmon sperm DNA
- Sortase reaction buffer: 50 mM Tris-HCl, 150 mM NaCl, pH 7.5, supplemented with 10 mM CaCl_2_

### PCA, PCD and Trolox

- 40× PCA was prepared by mixing 154 mg PCA in 10 ml water and NaOH, and adjusting the pH to 9.0.
- 100× PCD was prepared by adding 2.3 mg PCD to 3.3 ml of 100 mM Tris-HCl pH 8, 50 mM KCl, 1 mM EDTA, 50% glycerol
- 100× Trolox was prepared by first adding 100 mg Trolox to 430 μl of 100% methanol and 3.2 ml water, then adding 350 μl of 1 M NaOH and finally adding ∼480 μl of 1 M NaOH in 40 μl increments, shaking the solution each time, until the Trolox was completely dissolved.

#### Nanobody-DNA conjugation via a single cysteine

The conjugation of nanobodies to Combi-PAINT docking sites was performed as described previously^25^. First, the buffer was exchanged to 1× PBS + 5 mM EDTA, pH 7.0 using Amicon centrifugal filters (10k MWCO). Free cysteines were reacted with 20-fold molar excess of bifunctional EZ-Link Maleimide-PEG4-DBCO for 2-3 hours on ice. Unreacted linker was removed by buffer exchange to PBS using Amicon centrifugal filters. Azide-functionalized DNA was added with 3-to 5-molar excess to the DBCO-nanobody and reacted overnight at 4°C.

#### Nanobody-DNA conjugation via Sortase

Nanobodies carrying a C-terminal LPETG-HHHHHH tag were incubated with Sortase A 7M and GGG-DNA at a 0.25-fold and 3-fold molar ratio, respectively, in Sortase reaction buffer. Azide-DNA was reacted overnight at 4 °C with DBCO-PEG4-Gly-Gly-Gly to obtain GGG-DNA, which was then added to the sortase reaction. The reactions were then carried out at 25 °C for 1 h.

#### Nanobody purification

Unconjugated nanobody and free ssDNA were separated from DNA-conjugated nanobodies by anion exchange chromatography using an ÄKTA Pure liquid chromatography system equipped with a Resource Q 1 mL column. The concentrated DNA-conjugated nanobodies were adjusted to a concentration of 5-10 µM in 1× PBS containing 50% glycerol and 0.02% NaN_3_ and stored at -20 °C for 1-6 months or at -80 °C for >6 months.

#### Cloning

For transient transfection, gBlocks containing the genes of interest were ordered from IDT and cloned into a pcDNA3.1(+) mammalian expression vector (Thermo Fisher Scientific, cat. V79020) using Gibson assembly (NEB, E2611S).

#### DNA origami self-assembly

All DNA origami structures were designed in Picasso Design^11^. Self-assembly of DNA origami was accomplished in a one-pot reaction mix with a total volume of 50 μl, consisting of 10 nM scaffold strand (for sequence, see **Supplementary Data 1**), 100 nM folding staples (**Supplementary Data 2**), 500 nM biotinylated staples (**Supplementary Data 2**) and 1 μM staple strands with docking site extensions (**Supplementary Data 2**) in folding buffer. The reaction mix was then subjected to a thermal annealing ramp using a thermocycler. First, it was incubated at 80 °C for 5 min, cooled using a temperature gradient from 60 to 4 °C in steps of 1 °C per 3.21 min, and finally held at 4 °C.

#### DNA origami purification

DNA origami structures were purified via ultrafiltration using Amicon Ultra Centrifugal Filters with a 100-kDa molecular weight cutoff as previously described^26^. Folded origami was brought to 500 μl with FoB5 buffer and spun for 3.5 min at 10,000 *g*. This process was repeated twice. Purified DNA origami structures were recovered into a new tube by centrifugation for 5 min at 5,000 *g*. Purified DNA origami were stored at -20 °C in DNA LoBind tubes (Eppendorf, 0030108035).

#### DNA origami sample preparation

For sample preparation, a bottomless six-channel slide (ibidi, 80608) was attached to a coverslip. First, 200 μl of biotin-labeled BSA (1 mg/ml, dissolved in buffer A) was flushed into the chamber and incubated for 2 min. The chamber was then washed with 200 μl of buffer A. 200 μl neutravidin (0.1 mg/ml, dissolved in buffer A) was then flushed into the chamber and allowed to bind for 2 min. After washing with 200 μl of buffer A and subsequently with 200 μl of buffer B, 50 μl of biotin-labeled DNA structures (∼200 pM, roughly equal parts R1-R6, D1-D15 and T1-T20 20 nm grid structures where applicable) in buffer B was flushed into the chamber and incubated for 6 min. After DNA origami incubation the chamber was washed 3× with 200 μl of buffer B.

#### DNA origami imaging

For each round of Exchange-PAINT, 200 μl of the imager solution with 1xPCA, 1xPCD and 1xTrolox in buffer B was flushed into the chamber. Between imaging rounds, the sample was washed with 2 ml buffer B until no residual signal from the previous imager solution was detected. Then, the next imager solution was introduced. Combi-PAINT rounds were acquired in ascending order (R1*, R2*, … R6*). Imager sequences are listed in **Supplementary Table 2** and acquisition parameters (imager concentrations, number of frames acquired) are listed in **Supplementary Table 3**.

#### Cell culture

All cells were kept at 37°C, 5% CO_2_ and passaged every 2-3 days with 0.05% Trypsin-EDTA (Thermo Fisher Scientific, 25300-054). CHO-K1 cells were cultured in F12-K Nutrient Mix (Thermo Fisher Scientific, 21127022) supplemented with 10% FBS. Approximately 10,000 cells per square centimeter were seeded on eight well high glass bottom ibidi chambers (80807). A549 EGFR-GFP cells were cultured in DMEM medium (Thermo Fisher Scientific, 61965026) supplemented with 10% FBS (Thermo Fisher Scientific, 10500-064). Approximately 10,000 cells per square centimeter were seeded on eight well high glass bottom ibidi chambers.

#### Transient transfection of CHO-K1 cells

CHO-K1 cells were transfected with a pcDNA3.1 plasmid containing ALFA-Cd86-mEGFP using a Lipofectamine 3000 Kit (Thermo Fisher Scientific, L3000015) according to the manufacturer’s instructions. 8 wells of an ibidi high well slide were transfected with 2000 ng DNA in total, using Opti-MEM (Thermo Fisher scientific, 31985070), Lipofectamine 3000 and P3000 reagent. Cells were kept in the incubator for 16 to 24 hours before fixation.

#### Generation of pbCHO-K1 CD86-GFP

To generate doxycycline-inducible CD86-GFP stable cell lines, CHO-K1 cells were transfected in a six-well plate with 0.8 μg of the donor plasmid, 0.8 μg PiggyBac transposase vector^27^, and 0.8 μg reverse tetracycline transactivator plasmid^28^ using Lipofectamine3000 (Thermo Fisher Scientific, L3000001) according to the manufacturer’s instructions. After 48 hours, cells were plated at 40% confluency in a T25 flask, followed by selection with 1 μg/ml puromycin (Thermo Fisher Scientific, cat. A1113802) for six days. To induce transgene expression, 0.5 μg/ml doxycycline (Merck, cat. D9891) was added to the medium for at least 24 hours. GFP-positive cells were enriched using fluorescence-activated cell sorting on a CytoFLEX SRT (Beckman Coulter) and cells were maintained in F-12K medium (Thermo Fisher Scientific, cat. 21127030), supplemented with 10% fetal bovine serum and 1 μg/ml puromycin.

#### Cell fixation and staining

CHO-K1 Cd86-mEGFP (transient) were fixed with 4% pre-warmed (37°C) PFA for 20 minutes at RT and washed three times with 1xPBS before adding 0.1% Triton-X for 5 minutes at RT. After washing three times with 1xPBS the cells were blocked for 90 minutes with blocking buffer. 25nM anti-GFP nanobodies conjugated to the Combi-PAINT sequence of choice, in blocking buffer, were incubated overnight at 4°C. Before imaging, the cells were washed three times with 1xPBS, post-fixed with 4% pre-warmed (37°C) PFA for 10 minutes, washed three times with 1xPBS, quenched with 0.2M NH4Cl for 5 minutes, and washed three times with 1xPBS. Gold nanoparticles (diluted 1:2 in 1xPBS) were incubated for 7 minutes, and the cells were washed three times with 1xPBS.

A549-EGFR-GFP cells for Combi-PAINT characterization (**Fig. 3**) were fixed with 4% pre-warmed (37°C) PFA for 15 minutes at RT, washed three times with 1xPBS, permeabilized with 0.125% Triton-X for 2 minutes at RT, washed three times with 1xPBS, and blocked for 90 minutes with blocking buffer. 25nM of each anti-GFP nanobody conjugated to various Combi-PAINT sequences (e.g. D5, D9, D12, D14, D15) were incubated simultaneously in blocking buffer overnight at 4°C. Before imaging, the cells were washed three times with 1xPBS, post-fixed with pre-warmed (37°C) 4% PFA and 0.2% GA for 10 minutes, washed three times with 1xPBS, quenched with 0.2M NH4Cl for 5 minutes, and washed three times with 1xPBS. Gold nanoparticles (diluted 1:2 in 1xPBS) were incubated for 7 minutes, and the cells were washed three times with 1xPBS. pbCHO-K1 CD86 cells were prepared the same as transiently transfected CHO-K1, except that rather than one nanobody species, 25nM of each anti-GFP nanobody conjugated to various Combi-PAINT sequences (e.g. D5, D9, D12, D14, D15) were incubated simultaneously in blocking buffer overnight at 4°C.

#### CHO-K1 CD86, pbCHO-K1 CD86, and A549-EGFR-GFP imaging

For each round of Exchange-PAINT, 300 μl of the imager solution with 1xTolox in buffer C was added to the well. Between imaging rounds, the well was washed with 2-3 ml of PBS until no residual signal was detected. Then, the next imager solution was introduced. Combi-PAINT rounds were acquired in ascending order (R1*, R2*, … R6*). Imager concentrations and number of frames acquired are listed in **Supplementary Table 3**.

#### Microscope setup

Fluorescence imaging was carried out on an inverted microscope (Nikon Instruments, Eclipse Ti2) with the Perfect Focus System, applying an objective-type TIRF configuration equipped with an oil-immersion objective (Nikon Instruments, Apo SR HP TIRF ×100, NA 1.49, Oil). 488 nm (Toptica iBeam smart, 200 mW) and 560 nm (MPB Communications 2RU-VFL-P-1000-560-B1R, 1 W) lasers were used for excitation and coupled into the microscope via a Nikon manual TIRF module. The laser beams were passed through cleanup filters (Chroma Technology, ZET488/10x for 488 nm excitation and ZET561/10x for 560 nm excitation) and coupled into the microscope objective using a beam splitter (Chroma Technology, ZT488rdc-UF2 for 488 nm excitation and ZT561rdc-UF2 for 560 nm excitation). Fluorescence was spectrally filtered with an emission filter (Chroma Technology, ET525/50m and ET500lp for 488 nm excitation, and ET600/50m and ET575lp for 560 nm excitation) and imaged on an sCMOS camera (Hamamatsu, ORCA-Fusion BT) without further magnification, resulting in an effective pixel size of 130 nm (after 2×2 binning). The central 1152×1152 pixels (576×576 after binning) of the camera were used as the region of interest, and the scan mode was set to “ultra quiet scan”. Raw microscopy data was acquired using μManager (Version 2.0.1)^29^. TIR illumination was used for all acquisitions. The power density was determined as previously reported^21^ and set to 5 W/cm^2^ for 488 nm GFP screening and 150 W/cm^2^ for 560 nm DNA-PAINT imaging.

#### Image rendering

All microscopy images were rendered in Picasso Render^11^ using “individual localization precision, iso” as display setting, which is based on Equation 6 from Mortensen et al.^30^.

#### Combi-PAINT analysis

A schematic of the Combi-PAINT analysis workflow, including the decoding step, is illustrated in **Extended Data Figure 1**. Briefly, localizations from individual imaging rounds (**Extended Data Figure 1a**) were combined into a single file and clustered using a radial clustering algorithm (**Extended Data Figure 1b**) as described before^7^. Thus, localizations from all underlying imaging rounds contribute to clustering and identification, leading to sampling equivalent to standard DNA-PAINT. Localizations in sequential frames were identified as DNA-PAINT binding events (**Extended Data Figure 1c**, left). Next, each cluster was assigned an identity based on the channels in which the constituent binding events occurred, e.g., binding events in R1 and R2 channels led to the D1 identity assignment (**Extended Data Figure 1c**, right, **Supplementary Note 3**, and **Supplementary Figure 3**). A minimum number of binding events per cluster and a minimum number of binding events per channel were imposed to filter “sticking”, unspecific, events. Furthermore, the kinetics of all underlying channels were analyzed and filtered as described in **Supplementary Note 2**. The final Combi-PAINT image is rendered by merging the multi-base-channel localizations into the corresponding decoded combinatorial channel (**Extended Data Figure 1d**). All custom scripts used for Combi-PAINT were written in Python and are available at https://github.com/jungmannlab/combi-paint.

#### Combi-PAINT analysis of DNA origami

To determine the ground-truth ID of each DNA origami, the localizations coming from all 12 sites of the origami were merged. A ratio of approx. 20:1 was obtained between the specific and unspecific localizations, making the ID retrieval of each DNA origami robust and reliable. After the ID of each DNA origami was determined, the standard Combi-PAINT analysis was applied. The retrieved decoded IDs of single binding sites were compared with the ground-truth ID of the DNA origami.

#### Detection efficiency and decoding accuracy

##### Decoding Accuracy

Percentage of correctly decoded binding sites out of the space of sequences encoded in the sample (e.g., R, D, and T sequences for Figure 2, D sequences only for Figures 3 and 4).

##### Detection Efficiency

Percentage of decoded binding sites that belong to the set of encoded sequences in the sample out of the total number of decoded binding sites (including binding sites that are decoded as a sequence outside the encoded set). For example, for Figures 3 and 4, it is the number of decoded D sequences over the total number of decoded sequences (including the falsely decoded R and T sequences).

## Acknowledgments

We thank M. K. Steen-Mueller, U. Mueller and M. Joseph for proofreading the manuscript. P.R.S., I.P., R.K., L.H., M.H., and S.C.M.R. acknowledge support from the International Max Planck Research School for Molecules of Life (IMPRS-ML). L.A.M. acknowledges the postdoctoral fellowship from the European Union’s Horizon 20212022 research and innovation program under Marie Skłodowska-Curie grant agreement no. 101065980.

## Author information

P.R.S. and L.A.M. designed and performed experiments, developed analysis methods, and analyzed data. I.P., L.H., C.S., J.K., and M.H. prepared samples. R.K., S.C.M.R., and H.G. contributed to data analysis. P.R.S., L.A.M., and R.J. interpreted the data and wrote the manuscript with input from all authors. L.A.M. and R.J. conceived and supervised the project. P.R.S. and L.A.M. contributed equally.

## Ethic declarations

The authors declare no competing interests.

## References

1 Mund, A. et al. Deep Visual Proteomics defines single-cell identity and heterogeneity. Nat Biotechnol 40, 1231–1240 (2022). 10.1038/s41587-022-01302-5

2 Giesen, C. et al. Highly multiplexed imaging of tumor tissues with subcellular resolution by mass cytometry. Nat Methods 11, 417–422 (2014). 10.1038/nmeth.2869

3 Angelo, M. et al. Multiplexed ion beam imaging of human breast tumors. Nat Med 20, 436–442 (2014). 10.1038/nm.3488

4 Goltsev, Y. et al. Deep Profiling of Mouse Splenic Architecture with CODEX Multiplexed Imaging. Cell 174, 968–981 e915 (2018). 10.1016/j.cell.2018.07.010

5 Sahl, S. J., Hell, S. W. & Jakobs, S. Fluorescence nanoscopy in cell biology. Nat Rev Mol Cell Biol 18, 685–701 (2017). 10.1038/nrm.2017.71

6 Balzarotti, F. et al. Nanometer resolution imaging and tracking of fluorescent molecules with minimal photon fluxes. Science 355, 606–612 (2017). 10.1126/science.aak9913

7 Reinhardt, S. C. M. et al. Angstrom-resolution fluorescence microscopy. Nature 617, 711–716 (2023). 10.1038/s41586-023-05925-9

8 Sahl, S. J. et al. Direct optical measurement of intramolecular distances with angstrom precision. Science 386, 180–187 (2024). 10.1126/science.adj7368

9 Masullo, L. A. et al. Spatial and stoichiometric in situ analysis of biomolecular oligomerization at single-protein resolution. Nat Commun 16, 4202 (2025). 10.1038/s41467-025-59500-z

10 Jungmann, R. et al. Single-molecule kinetics and super-resolution microscopy by fluorescence imaging of transient binding on DNA origami. Nano Lett 10, 4756–4761 (2010). 10.1021/nl103427w

11 Schnitzbauer, J., Strauss, M. T., Schlichthaerle, T., Schueder, F. & Jungmann, R. Super-resolution microscopy with DNA-PAINT. Nat Protoc 12, 1198–1228 (2017). 10.1038/nprot.2017.024

12 Jungmann, R. et al. Multiplexed 3D cellular super-resolution imaging with DNA-PAINT and Exchange-PAINT. Nat Methods 11, 313–318 (2014). 10.1038/nmeth.2835

13 Unterauer, E. M. et al. Spatial proteomics in neurons at single-protein resolution. Cell 187, 1785–1800 e1716 (2024). 10.1016/j.cell.2024.02.045

14 Schueder, F. et al. Unraveling cellular complexity with transient adapters in highly multiplexed super-resolution imaging. Cell 187, 1769–1784 e1718 (2024). 10.1016/j.cell.2024.02.033

15 Chen, K. H., Boettiger, A. N., Moffitt, J. R., Wang, S. & Zhuang, X. RNA imaging. Spatially resolved, highly multiplexed RNA profiling in single cells. Science 348, aaa6090 (2015). 10.1126/science.aaa6090

16 Lubeck, E. & Cai, L. Single-cell systems biology by super-resolution imaging and combinatorial labeling. Nat Methods 9, 743–748 (2012). 10.1038/nmeth.2069

17 Ben-Uri, R. et al. High-dimensional imaging using combinatorial channel multiplexing and deep learning. Nat Biotechnol (2025). 10.1038/s41587-025-02585-0

18 Strauss, S. & Jungmann, R. Up to 100-fold speed-up and multiplexing in optimized DNA-PAINT. Nat Methods 17, 789–791 (2020). 10.1038/s41592-020-0869-x

19 Rothemund, P. W. Folding DNA to create nanoscale shapes and patterns. Nature 440, 297–302 (2006). 10.1038/nature04586

20 Bennett, N. R. et al. Improving de novo protein binder design with deep learning. Nat Commun 14, 2625 (2023). 10.1038/s41467-023-38328-5

21 Steen, P. R. et al. The DNA-PAINT palette: a comprehensive performance analysis of fluorescent dyes. Nat Methods 21, 1755–1762 (2024). 10.1038/s41592-024-02374-8

22 Barentine, A. E. S. et al. An integrated platform for high-throughput nanoscopy. Nat Biotechnol 41, 1549–1556 (2023). 10.1038/s41587-023-01702-1

23 Geertsema, H. J. et al. Left-handed DNA-PAINT for improved super-resolution imaging in the nucleus. Nat Biotechnol 39, 551–554 (2021). 10.1038/s41587-020-00753-y

24 Banerjee, A., Anand, M., Srivastava, M., Vidwath, V. S. & Ganji, M. High-Speed Multiplexed DNA-PAINT Imaging of Nuclear Organization Using an Expanded Sequence Repertoire. bioRxiv, 2024.2012.2021.629871 (2025). 10.1101/2024.12.21.629871

25 Sograte-Idrissi, S. et al. Nanobody Detection of Standard Fluorescent Proteins Enables Multi-Target DNA-PAINT with High Resolution and Minimal Displacement Errors. Cells 8 (2019). 10.3390/cells8010048

26 Wagenbauer, K. F. et al. How We Make DNA Origami. Chembiochem 18, 1873–1885 (2017). 10.1002/cbic.201700377

27 Li, Z., Michael, I. P., Zhou, D., Nagy, A. & Rini, J. M. Simple piggyBac transposon-based mammalian cell expression system for inducible protein production. Proc Natl Acad Sci U S A 110, 5004–5009 (2013). 10.1073/pnas.1218620110

28 Yusa, K., Zhou, L., Li, M. A., Bradley, A. & Craig, N. L. A hyperactive piggyBac transposase for mammalian applications. Proc Natl Acad Sci U S A 108, 1531–1536 (2011). 10.1073/pnas.1008322108

29 Edelstein, A. D. et al. Advanced methods of microscope control using muManager software. J Biol Methods 1 (2014). 10.14440/jbm.2014.36

30 Mortensen, K. I., Churchman, L. S., Spudich, J. A. & Flyvbjerg, H. Optimized localization analysis for single-molecule tracking and super-resolution microscopy. Nat Methods 7, 377–381 (2010). 10.1038/nmeth.1447

