## Supplementary Information for "Combinatorial DNA-PAINT"

| | |
|---|---|
| **Supplementary Table 1** | **Combi-PAINT docking sequences** |
| **Supplementary Table 2** | **Imager sequences** |
| **Supplementary Table 3** | **Acquisition parameters** |
| **Supplementary Note 1** | **Relative acquisition speed** |
| **Supplementary Figure 1** | **Relative acquisition speed of various DNA-PAINT multiplexing schemes** |
| **Supplementary Note 2** | **Filtering** |
| **Supplementary Figure 2** | **Filtering** |
| **Supplementary Note 3** | **Identification of Combi-PAINT ID** |
| **Supplementary Figure 3** | **Detection efficiency and decoding accuracy depending on identification criteria** |

**Supplementary Table 1. Combi-PAINT docking sequences**

| Name | Base sequence names | Sequence |
|---|---|---|
| R1 | R1 | TCCTCCTCCTCCTCCTCCT |
| R2 | R2 | ACCACCACCACCACCACCA |
| R3 | R3 | CTCTCTCTCTCTCTCTCTC |
| R4 | R4 | ACACACACACACACACACA |
| R5 | R5 | CTTCTTCTTCTTCTTCTTC |
| R6 | R6 | AACAACAACAACAACAACAA |
| D1 | R1 – R2 | AAACCACCACCACCATCCTCCTCCTCCT |
| D2 | R1 – R3 | TCCTCTCTCTCTCTCCTCCTCCTCCT |
| D3 | R1 – R4 | CAACACACACACACATCCTCCTCCTCCT |
| D4 | R1 – R5 | CCCTTCTTCTTCTTCCTCCTCCTCCT |
| D5 | R1 – R6 | AAAACAACAACAACAATCCTCCTCCTCCT |
| D6 | R2 – R3 | TCCTCTCTCTCTCTCACCACCACCACCA |
| D7 | R2 – R4 | CAACACACACACACACCACCACCACCA |
| D8 | R2 – R5 | CCCTTCTTCTTCTTCACCACCACCACCA |
| D9 | R2 – R6 | AAAACAACAACAACAACCACCACCACCA |
| D10 | R3 – R4 | CAACACACACACACACTCTCTCTCTCTC |
| D11 | R3 – R5 | CCCTTCTTCTTCTTCTCTCTCTCTCTC |
| D12 | R3 – R6 | AAAACAACAACAACAACTCTCTCTCTCTC |
| D13 | R4 – R5 | CCCTTCTTCTTCTTCACACACACACACA |
| D14 | R4 – R6 | AAAACAACAACAACAACACACACACACACA |
| D15 | R5 – R6 | AAAACAACAACAACAACTTCTTCTTCTTC |
| T1 | R1 – R2 – R3 | TCCTCTCTCTCTCTCACCACCACCACCATCCTCCTCCTCCT |
| T2 | R1 – R2 – R4 | CAACACACACACACACCACCACCACCATCCTCCTCCTCCT |
| T3 | R1 – R2 – R5 | CCCTTCTTCTTCTTCACCACCACCACCATCCTCCTCCTCCT |
| T4 | R1 – R2 – R6 | AAAACAACAACAACAACCACCACCACCATCCTCCTCCTCCT |
| T5 | R1 – R3 – R4 | CAACACACACACACACTCTCTCTCTCTCCTCCTCCTCCT |
| T6 | R1 – R3 – R5 | CCCTTCTTCTTCTTCTCTCTCTCTCTCCTCCTCCTCCT |
| T7 | R1 – R3 – R6 | AAAACAACAACAACAACTCTCTCTCTCTCCTCCTCCTCCT |
| T8 | R1 – R4 – R5 | CCCTTCTTCTTCTTCACACACACACACATCCTCCTCCTCCT |
| T9 | R1 – R4 – R6 | AAAACAACAACAACAACACACACACACACATCCTCCTCCTCCT |
| T10 | R1 – R5 – R6 | AAAACAACAACAACAACTTCTTCTTCTTCCTCCTCCTCCT |
| T11 | R2 – R3 – R4 | CAACACACACACACACTCTCTCTCTCTCACCACCACCACCA |
| T12 | R2 – R3 – R5 | CCCTTCTTCTTCTTCTCTCTCTCTCTCACCACCACCACCA |
| T13 | R2 – R3 – R6 | AAAACAACAACAACAACTCTCTCTCTCTCACCACCACCACCA |
| T14 | R2 – R4 – R5 | CCCTTCTTCTTCTTCACACACACACACACCACCACCACCA |
| T15 | R2 – R4 – R6 | AAAACAACAACAACAACACACACACACACACCACCACCACCA |
| T16 | R2 – R5 – R6 | AAAACAACAACAACAACTTCTTCTTCTTCACCACCACCACCA |
| T17 | R3 – R4 – R5 | CCCTTCTTCTTCTTCACACACACACACACTCTCTCTCTCTC |
| T18 | R3 – R4 – R6 | AAAACAACAACAACAACACACACACACACACTCTCTCTCTCTC |
| T19 | R3 – R5 – R6 | AAAACAACAACAACAACTTCTTCTTCTTCTCTCTCTCTCTC |
| T20 | R4 – R5 – R6 | AAAACAACAACAACAACTTCTTCTTCTTCACACACACACACA |

**Supplementary Table 2. Imager sequences**

| Name | Sequence | 3'-Modification |
|---|---|---|
| R1* | AGGAGGA | Cy3B |
| R2* | TGGTGGT | Cy3B |
| R3* | GAGAGAG | Cy3B |
| R4* | TGTGTGT | Cy3B |
| R5* | GAAGAAG | Cy3B |
| R6* | TTGTTGTT | Cy3B |

**Supplementary Table 3. Acquisition parameters**

| Experiment | Imagers | Imager concentration [pM] | Frames |
|---|---|---|---|
| R, D, T DNA origami (Fig. 2) | R1* | 800 | 20000 |
| | R2* | 200 | 20000 |
| | R3* | 1200 | 20000 |
| | R4* | 500 | 20000 |
| | R5* | 2000 | 20000 |
| | R6* | 2000 | 20000 |
| R, D, T DNA origami (high speed) | R1* | 1620 | 2400 |
| | R2* | 580 | 2400 |
| | R3* | 1020 | 2400 |
| | R4* | 1450 | 2400 |
| | R5* | 4560 | 2400 |
| | R6* | 3080 | 2400 |
| D sequences in CHO-K1 (Fig. 3a-e) | R1* | 500 | 40000 |
| | R2* | 250 | 40000 |
| | R3* | 400-500 | 40000 |
| | R4* | 250-500 | 40000 |
| | R5* | 500 | 40000 |
| | R6* | 400-500 | 40000 |
| D sequences in A549 (Fig. 3f-i) | R1* | 1200 | 30000 |
| | R2* | 300 | 30000 |
| | R3* | 1200 | 30000 |
| | R4* | 500 | 30000 |
| | R5* | 2000 | 30000 |
| | R6* | 1000 | 30000 |
| D sequences in pbCHO-K1 (Fig. 3i) | R1* | 1000 | 30000 |
| | R2* | 200 | 30000 |
| | R3* | 1500 | 30000 |
| | R4* | 250 | 30000 |
| | R5* | 1500 | 30000 |
| | R6* | 1500 | 30000 |

The exposure time was 100 ms and the power density 150 W/cm$^2$ for all experiments.

**Supplementary Note 1: Relative acquisition speed**

Factors determining DNA-PAINT acquisition speed include imaging time per round, which is dependent on imager concentration and the time necessary to sufficiently sample targets, and the time between imaging rounds, i.e. to perform buffer exchange.

In **Supplementary Figure 1** we plotted relative acquisition times, i.e. total acquisition time ($T_{Total}$) divided by the time needed to acquire a single target ($T_{Target}$) multiplied by the number of targets ($n_{Targets}$), against the number of targets.

$$Relative\ acquisition\ speed = \frac{T_{Total}}{T_{Target} \times n_{Targets}}$$

A lower value indicates faster imaging since less time is needed per acquired target. Notably, due to sequential imaging and buffer exchange times between imaging rounds (and secondary strand hybridization for SUM-PAINT), Exchange-PAINT and SUM-PAINT experiments always take longer than the corresponding number of single experiments ($T_{rel} > 1$). Combi-PAINT, on the other hand, takes less time per target the more targets are acquired - down to approx. 15% the time per target compared to individual imaging rounds.

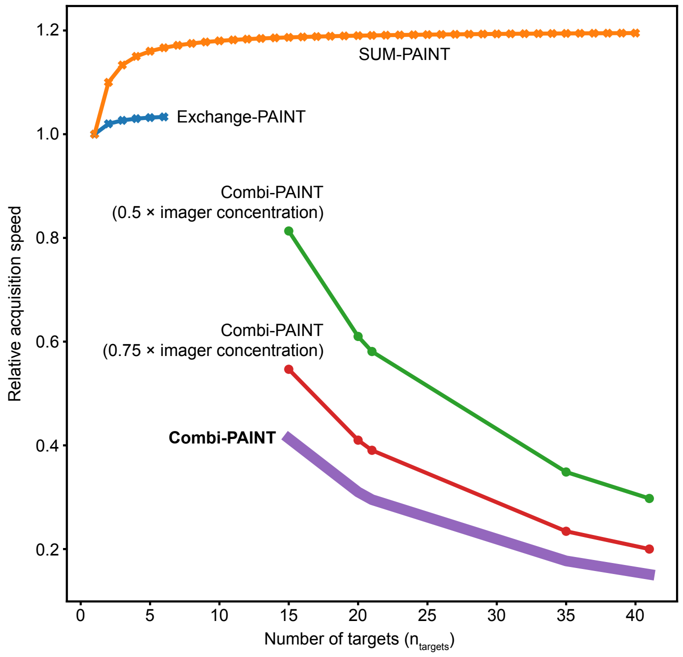


**Supplementary Figure 1 | Relative acquisition speed of various DNA-PAINT multiplexing schemes.** Dots indicate calculated values, solid lines are visual aids. Combi-PAINT line is highlighted as it is the optimal and theoretical maximum improvement.

**Supplementary Note 2: Filtering**

The standard deviation ($\sigma$) of the uniformly distributed times at which binding events occur ($t_{BE}$) is expected to be:

$$\sigma(t_{BE}) = \frac{T_{total}}{\sqrt{12}}$$

Experimentally, $\sigma(t_{BE})$ for all docking sites in a given measurement exhibit a certain variance. Clusters resulting from "sticking" events, however, generally feature unspecific binding events with a far lower standard deviation than genuine DNA-PAINT as "sticking" mostly occurs at a single time for a given location. We plotted $\sigma(t_{BE})$ from all clusters for R1 to R6 measurements with 30,000 frames (see **Supplementary Figure 2**) and applied a multi-component fit (red curve) featuring a Gaussian (blue curve) and third-degree polynomial (green curve) to account for the sticking, to isolate DNA-PAINT docking sites from likely unspecific sticking. The mean $\mu(t_{BE})$ = (8336.2 ± 831.7) frames across R1-R6 measurements corresponds well to the expected value of $\frac{30{,}000}{\sqrt{12}} \approx 8{,}660$ frames.

Using μ - 3×σ = 5841.1 frames as a threshold to delineate DNA-PAINT from non-specific sticking, we applied $0.2 \times n_{frames}$ = 6,000 (black vertical line) as a robust and straight-forward approximation for all measurements.

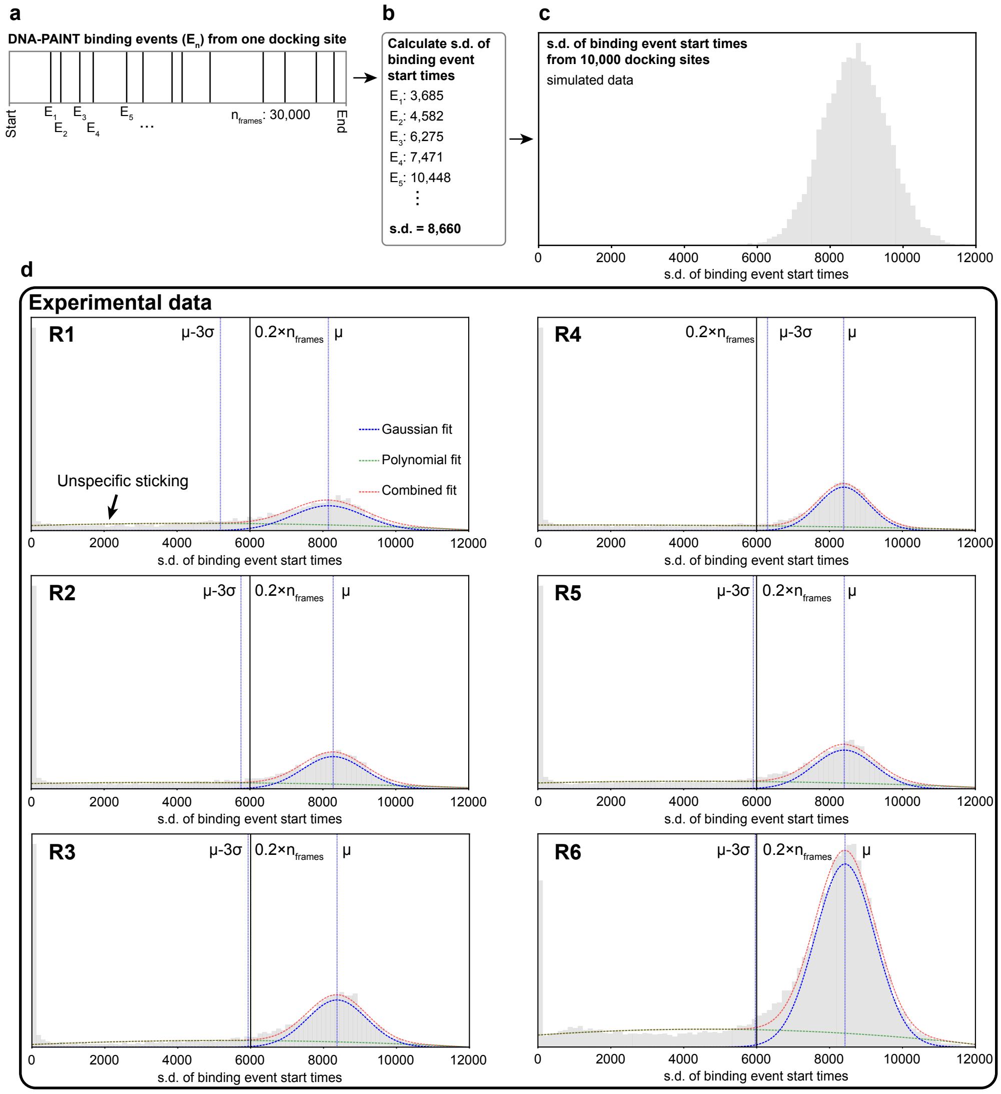

**Supplementary Figure 2 | Filtering. a,** DNA-PAINT binding events from one docking site. **b,** the standard deviation of the times at which binding events occur is calculated for each docking site. **c**, all standard deviations are plotted as a histogram. **d,** the experimental s.d. per docking site follow the expected distribution. Unspecific sticking is filtered by removing docking sites with s.d. below $0.2 \times n_{frames}$.

### Supplementary Note 3: Identification of Combi-PAINT ID

Clusters are assigned a Combi-PAINT ID if:

- There are X localizations or more
- There are Y binding events or more in total across all channels
- There are Z binding events or more per channel
- At least one channel passes the standard deviation filter (**see Supplementary Note 2**)

X and Y are determined by the degree of sampling in the experiment. Z was determined by screening Z = (1, 2, 3, 4) and selecting for best performance. We found that the total number of sites detected, DE, DA, and total number of correct sites are robust within this range, but optimal results are achieved for Z = 2 (see **Supplementary Figure 3**). We additionally examined the effect of mild variations in filtering cut-off and cluster radius, and see that the Combi-PAINT analysis is robust against small deviations from optimal parameters.

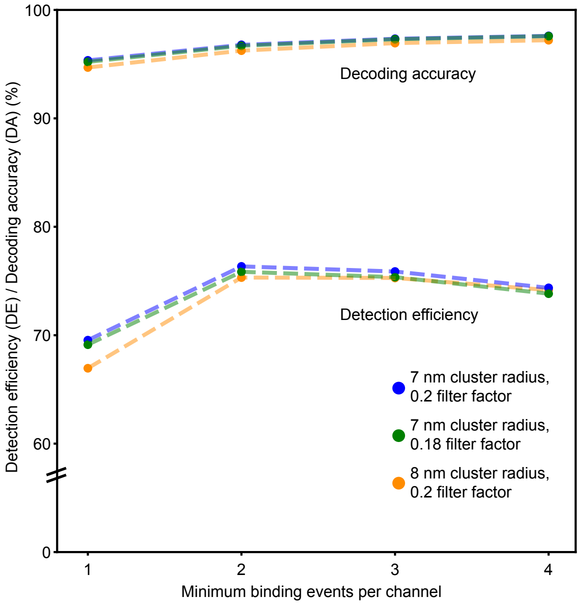


**Supplementary Figure 3 | Detection efficiency and decoding accuracy depending on identification criteria.**
